# Interleukin 11-induced microRNAs as functional mediators and circulating biomarkers of cardiac fibrosis

**DOI:** 10.1101/2023.10.11.561968

**Authors:** Roman Tikhomirov, Benedict Reilly-O’Donnell, Carla Lucarelli, Prashant Kumar Srivastava, Maryam Anwar, Chi Him Kendrick Yiu, Julia Dielesen, Santiago Nicolas Piella, Zoe Kwan, Germana Zaccagnini, Catherine Mansfield, Maddalena Tessari, Lorenzo Menicanti, Simona Greco, Przemysław Leszek, Giuseppe Faggian, Costanza Emanueli, Fabio Martelli, Julia Gorelik

**Author notes:** **Address of correspondence:** Julia Gorelik, Imperial College Hammersmith Hospital campus-ICTEM building-office 429; Du Cane Road, London W12 0HS, Fabio Martelli, Via Morandi 30, San Donato Milanese, Milan, 20097, Costanza Emanueli, Imperial College Hammersmith Hospital campus-ICTEM building-office 434; Du Cane Road, London W12 0HS.

## Abstract

**Background:** Cardiac fibrosis can be triggered by several pathologies, including ischemic heart disease and aortic stenosis (AS). Cardiac fibrosis is brought about by uncontrolled extracellular matrix (ECM) deposition by myofibroblasts. Interleukin-11 (IL-11) has been firmly demonstrated to be a major trigger of multi-organ fibrosis. However, the molecular mechanisms underpinning IL-11-induced fibrosis requires further characterisation. Recent studies indicate that microRNA (miRNA) dysregulation contributes to the pathogenesis of cardiac fibrosis and can be targeted therapeutically. In this study, we explored the hypothesis that miRNAs act as downstream effectors of IL-11-induced cardiac fibrosis. Moreover, we investigated the translational potential of IL-11-regulated miRNAs as circulating biomarkers of cardiac fibrosis in AS patients.

**Methods and Results:** Using computational approaches, we identified miRNA-497-5p and miRNA-27b-5p as potential new downstream profibrotic effectors of IL-11 in fibroblasts. We next confirmed that both miRNAs increased in healthy rat CF stimulated with IL-11 and in CF derived from post-infarction failing hearts. At the functional level, miRNA-497-5p and miRNA-27b-5p inhibition indirectly reduced the mRNA expression of collagen 1 (Col1a1). Conversely, transfection of CFs with mimics for each of the two miRNAs promoted fibroblast-to-myofibroblast transition and increased Col1a1 levels. We provided evidences that miRNA-27b-5p and miRNA-497-5p converge to promote hypoxia-inducible factor 1 signalling, by targeting its regulator EGLN (PHD) family members. The clinical relevance of our findings was confirmed using left ventricle (LV) specimens obtained from surgical patients with AS. The miRNA-27b-5p and miRNA-497-5p measured in the LV, peripheral plasma and plasma extracellular vesicles correlated with the severity of LV fibrosis, indicating these miRNAs’ potential as new circulating biomarkers of cardiac fibrosis.

**Conclusions:** In this study, we have newly identified the potential value of miRNA-27b-5p and miRNA-497-5p as actionable biomarkers of the profibrotic response to IL-11 in the heart. Future studies should validate the translational potential of the miRNAs as new clinical biomarkers and therapeutic targets.

## Introduction

Cardiac fibrosis is a pathological condition that accompanies various heart diseases [1]. It impairs heart function by extensive deposition of ECM proteins, especially collagens, into the myocardium [2]. ECM remodelling increases heart stiffness, driving heart failure (HF) [3]. CFs are the main source of ECM proteins. Activated fibroblasts or myofibroblasts can be characterised by α-SMA stress fibres and increased expression of pro-fibrotic factors. They play a key role in cardiac fibrosis by participating in ECM re-organization and potentiating further fibroblast activation [4]. CFs can be activated by transforming growth factor β1 (TGFβ1), which induces fibrosis-related gene transcription through the SMAD cascade [5]. Schafer et al. recently revealed that exogenous TGFβ1 supplementation increased the levels of IL-11 in primary CFs [6]. Further experiments showed that the activation of the IL-11ra receptor for IL-11 provokes a strong pro-fibrotic state in CFs through extracellular regulated kinase (ERK) phosphorylation, and transcriptional regulating collagens, α-SMA and autocrine IL-11 production. Accordingly, *IL-11ra* knock out mice were protected from developing fibrosis [6]. Interestingly, IL-11 was found to be increased in distinct cardiac and extra-cardiac conditions linked to ECM remodelling, such as myocardial infarction (MI) [7], pressure-overload [8] and idiopathic lung fibrosis [9–10]. The relevance of IL-11 for human medicine is further suggested by previous work on clinical samples. Plasma levels of IL-11 were increased in congestive HF patients, when compared with less severe cardiac patients [11]. Moreover, increased IL-11 was detected in the aortic tissue and plasma of patients with acute thoracic aortic dissection [12]. Taken together, these studies candidate IL-11 to the dual role of therapeutic target and biomarker of cardiac fibrosis. Notwithstanding, the downstream effectors and non-canonical signalling pathways triggered by IL-11 in CFs remains largely unexplored [13]. One class of molecules which could regulate the action of IL-11 in CFs are microRNAs (miRNAs), the most studied class of small non-coding RNAs. The canonical action of individual miRNA consists of post-transcriptional repression of gene expression by their binding to pool specific messenger RNA (mRNAs) to induce mRNA degradation or translational inhibition [14]. Several miRNAs have already been shown to be relevant for human cardiovascular disease and cardiac fibrosis [14–17] and to mediate the fibrotic response triggered by TGFβ1 [18–20]. Moreover, miRNAs have been proposed to have value as circulating biomarkers of organ fibrosis [21], especially when circulating in small sEVs, also referred to as “exosomes” [22]. The involvement of miRNAs in the cardiac fibrotic response to IL-11 remains unexplored.

AS is caused by the progressive calcification and reduced mobility of the aortic valve leaflets, leading to pressure overload on the LV, hypertrophic and fibrotic remodelling and consequent HF. AS is the most frequent heart valve disease in the western world, with more than 2% of patients over 60 years suffering from this condition [23]. Treatments for AS include surgical aortic valve replacement (SAVR) or transcatheter aortic valve implantation (TAVI). If left untreated, the mortality rate associated with AS reaches 50% at one year after symptoms onset [24]. Current guidelines recommend treatment for AS in symptomatic patients or in case of LV impairment. However, symptoms are challenging to assess in this cohort of patients. Recent imaging studies have demonstrated that even a minor LV impairment is associated with worse outcomes after SAVR or TAVI [25–26]; and LV fibrosis is associated with increased long-term mortality despite treatment [27]. Therefore, imaging and laboratory biomarkers enabling to estimate the of fibrosis in the LV would crucially aid in planning the timing for intervention in these patients to prevent a subtle yet significant myocardial damage and maladaptive remodelling, ultimately improving outcomes. Cardiac magnetic resonance (CMR) with late gadolinium enhancement (LGE) is the gold standard to study cardiac fibrosis. Despite CMR being an extremely useful technique, there are several limitations intrinsic to this technique [28]. Identifying compelling laboratory biomarkers associated with cardiac fibrosis has the potential to improve our capacity to determine the optimal time of intervention in patients with severe AS but without overt symptoms. Circulating miRNAs have the potential to meet this aim.

In this study, we have combined work *in silico*, with cell and molecular biology and analyses of clinical samples from a highly selected population of patients with severe AS. This has led us to newly identify and assess the potential value of miRNA-27b-5p and miRNA-497-5p as novel therapeutic targets and circulating indicators of cardiac fibrosis.

## Methods

For detailed methods, see the Supplemental Appendix.

### Ethics regulation of work on human and animal samples

Studies on human tissue followed the principles outlined in the Helsinki Declaration and the Italian and Polish laws and guidelines and were authorised by local Ethics Committees (for Italy: protocol #2438, 27/01/2009 and CE#85/int/2016 9/6/2016; for Poland protocol # IK-NPIA-0021-14/1426/18). Studies on human organotypic culture was approved by the NHS committee (REC reference 19/SC/0257; IRAS project ID:264059). Animal work was developed entirely in the United Kingdom (UK) with established standards for the care of animal subjects written by the UK Home Office (ASPA1986 Amendments Regulations 2012) incorporating the EU directive 2010/63/EU.

### Cell culture

Human CFs (hCFs) were obtained from organotypic culture of LV tissue collected from healthy donor heart rejected for transplantation (N = 1, n = 4). Rat CFs (rCFs) were isolated following enzymatic digestion of the LV of healthy rats or rats at 16 weeks post-MI HF (N = 3-4). HF was confirmed by measuring fraction shortening and ejection fraction through echocardiogram (Supplementary Figure 1).

Culturing protocols result in high purity of fibroblasts, which was confirmed by the PDGFRα and vimentin staining (Supplementary Figure 2). In the experimental conditions, vimentin was used as a cytosolic stain of fibroblasts and myofibroblasts. α-SMA was used as a myofibroblast specific stain in our cultures. Both hCFs and rCFs were cultured in high-glucose (4500 mg/l) Dulbecco’s Modified Eagle Medium no longer than 3 weeks, passage 3-5 [29], cells were seeded before following experiment in 12 well plate at 70% confluence.

### Prediction of miRNAs involved in regulation of cardiac fibrosis, miRNA targets prediction

A graphical abstract for the protocol is described in supplementary figure 3. To predict miRNAs involved in the regulation of cardiac fibrosis we (i) first identified genes in IL-11 (ERK) and canonical TGFβ1 pathways only using the KEGG pathway database. Although TGFβ1 induces IL-11 production, the downstream signalling via ERK differs from the canonical SMAD cascade [30]. Highly expressed miRNAs in CFs (top 25 %) were identified by (ii) using publicly available dataset downloaded from Gene Expression Omnibus ID: GSE76175. From the remaining miRNAs in the same dataset, we extracted(iii) miRNAs which have evidence for target regulation [31] within a list of genes identified in the first step. miRNAs from step (ii) and (iii) were analysed *in silico* for binding affinity with gene targets obtained from the KEGG pathway database (i) by intersection of three well known prediction tools: TargetScan, miRWalk and miRcode [32–34]. miRNAs were considered for further analysis if they were predicted to bind by all three tools, were localized in 3’-UTR of the target gene [35] and were conserved between human and rat. The resulting miRNAs were shortlisted based on their number of targets within the IL-11 (ERK) and TGFβ1 signalling pathways (№ of targets ≥ 3).

### RNAseq on human cardiac fibroblasts stimulated with IL-11

RNAseq was designed to identify genes in CFs regulated by IL-11 stimulation and was used for the enrichment analysis. Total RNA was isolated with Trizol from human dilated cardiomyopathy CFs (N = 3) treated with IL-11 (5ng/µl for 24 hours) and from non-stimulated fibroblasts (N = 3). Isolation purity war proved by a comparison of the absorbance of nucleic acids at 260 nm with following wavelengths: 230 nm and 280 nm. Ribosomal RNA was depleted. The Imperial BRC Genomics facility performed RNAseq. Further differential gene expression analysis was performed using the EdgeR [36] tool implemented in R as described previously [37].

### Regulation of new molecular targets in IL-11 signalling pathway by miRNA-27b-5p and miRNA-497-5p

To identify new molecular targets in IL-11 signalling pathway we applied the approach above. The resulting targets of miRNA-27b-5p (№ of targets = 596) and miRNA-497-5p (№ of targets = 140) were further filtered by their expression in CFs after IL-11 stimulation. Next, over-representation analysis for resulting targets was performed by WebGestalt [38] with default settings. The most enriched pathway was determined by p-values (p-value <0.05). Targets from this pathway were used to plot the gene network in Cytoscape [39].

### Immunostaining of cardiac fibroblasts

hCFs and rCFs were stained with primary antibodies overnight: PDGFRα rabbit monoclonal antibody (ab203491), α-SMA mouse monoclonal DAKO antibody (M0851) and vimentin chicken polyclonal antibody from Invitrogen (PA1-16759).

### 3’UTR luciferase assays to demonstrate miRNA direct binding to the predicted target genes

To demonstrate the direct binding and expressional control exerted by miR-27b and miR-497 on target genes emerging from the aforementioned bioinformatic analyses, we adapted a previously described protocol [40] HEK293FT cells were seeded in 96-well plates and transfected with WT, MUT target constructs or vector in combination with miRNA control (Scramble - GeneCopoeiaTM CmiR0001-MR04) or with Homo sapiens miR-27b stem-loop (GeneCopoeiaTM HmiR0145-MR04) or with Homo sapiens miR-497 stem loop (GeneCopoeiaTM HmiR-0271-MR04) and with pRL-null renilla luciferase (N = 4 for each combination).

### Clinical samples

Table 1 summarises the characteristics of patients and healthy donors included in this study: 1) patients with severe high gradient AS pstients undergoing elective surgical aortic valve replacement (SAVR) who donated blood and heart tissues; 2) healthy volunteers who donated blood (controls). Moreover, control healthy left ventricle samples were obtained from unused transplantation donor hearts. The AS patients were prospectively recruited (2018 to 2021) at the Policlinico G.B. Rossi, Verona, Italy. The patients selected for the present study had symptomatic severe, high-gradient AS confirmed on echocardiogram mean gradient ≥ 40 mmHg, peak velocity ≥ 4.0 m/s and valve area ≤ 1.0 cm^2^ (or ≤ 0.6 cm^2^/m^2^), trileaflet aortic valve and preserved left ventricular ejection fraction (LVEF) ≥ 55 %. Exclusion criteria were: (1) bicuspid anatomy, (2) low flow-low gradient AS, (3) history of coronary artery disease or significant coronary artery disease with indication to surgical revascularisation concomitant to SAVR, (4) impaired left ventricular function, (5) history of arrhythmia including atrial fibrillation, (6) chronic kidney disease and liver dysfunction. Patients with significantly hypertrophic septum underwent concomitant septal myectomy. Blood plasma and LV samples (frozen or formalin fixed) were collected. Detailed characteristics and comorbidities of the AS patients are presented in Table 2. Control blood donors were recruited at IRCCS Policlinico San Donato, Milan, Italy. Donor hearts were provided by the Department of Heart Failure and Transplantology, Cardinal Stefan Wyszyński Institute of Cardiology, Warsaw, Poland.

### Histology and morphometric analysis of the human samples

Heart Samples were fixed and embedded in paraffin according to standard procedures. ECM was stained in tissue histology sections of AS (N = 11) and control samples (N = 10) using a Picro Sirius Red Stain Kit. For immunofluorescence staining the following antibodies were used: anti-collagen 1 (ab34710 Abcam Cambridge, UK) (1:200 v/v dilution in 5% BSA in PBS tween 0.05%) and anti-alpha sarcomeric actin clone 5C5 (#2172, Sigma-Aldrich, Merck Darmstadt, Germany) (1:500 v/v dilution in 5% BSA in PBS tween 0.05%). Collagen was quantified according to the protocol described by Ying Chen and colleagues [41].

### Plasma extracellular vesicle analysis

Citrate plasma was prepared from peripheral blood of AS (N = 23) and control donors (N = 10) and stored at −80 ^0^C freezers. Plasma sEVs were isolated by size exclusion chromatography, using the Exo-spin blood kit following the manufacturer’s protocol (CELL Guidance systems). The quality of the sEVs preparation was validated using transmission electron microscopy with 2% uranyl acetate staining (transmission electron microscope TALOS L120C ThermoScientific at 120 kv).

### RNA isolation and reverse transcription polymerase chain reaction

Total RNA isolation from cell cultures and LV biopsies was performed by Trizol (ThermoFisher Scientific) following the manufacturer’s protocol. GoScript Reverse Transcriptase (Promega) was used to generate cDNAs. Genes analyzed in this study are indicated in supplementary table 1. miRNAs and small RNAs were isolated from plasma and plasma-EVs of AS patients and healthy donors by using Nucleospin miRNA Plasma kit (Macherey Nagel), after the addition of cel-miR-39 spike-in for normalization (5 pmol/µl) (ThermoFisher Scientific). MicroRNAs and U6 small nuclear RNA (supplementary table 2) were polyadenylated and converted into cDNA by using TaqManTM MicroRNA Reverse Transcription (RT) kit (Thermo Fisher Scientific). Amplification was visualized by StepOnePlus™ Real-Time PCR System (ThermoFisher Scientific).

### Statistics

Statistical analysis was performed with Origin 8 Pro and Rstudio. The tests performed were indicated in the figure legends. Normal distribution of the data was determined by a Shapiro-Wilk test. For two groups comparison Mann-Whitney test was applied. For multiple experimental groups comparison One-Way ANOVA with Tukey’s post-hoc test was used. Correlations with miRNA-27b-5p, miRNA-497-5p and EGLN1 expression levels were quantified with Pearson’s correlation test. Receiver operative characteristics (ROC) curves were used to assess the diagnostic accuracy of miRNAs in distinguish healthy donors of plasma and AS patients. N= number of patients/ animals, n= number of technical repeats. Data were presented as Mean ± SEM. P values <0.05 were considered statistically significant.

## Results

### Cardiac fibroblasts are activated upon *in vitro* treatment with IL-11 and in the post-MI failing heart

Using α-SMA as a marker of myofibroblasts in a cell culture with high fibroblast purity, we first confirmed that IL-11 administration in vitro promoted differentiation of rCFs in myofibroblasts, as well as post-MI HF condition (Figure 1 A). These data were corroborated by IHC analysis, showing increased abundance of α-SMA-positive CFs with recognizable stress fibres (Figure 1 B). In parallel, increased expression of α-SMA (Figure 1 C) was observed in rCFs treated with IL-11 (PBS) or derived from post-MI failing rat hearts (sham-operated hearts). Moreover, increased Col1a1 mRNA levels were detected in CFs stimulated with IL-11 in vitro (control: PBS) and in CFs prepared from the rat failing hearts (control: rat healthy hearts) (Figure 1 D) but not Col3a1 (Figure 1E).

**Figure 1.**
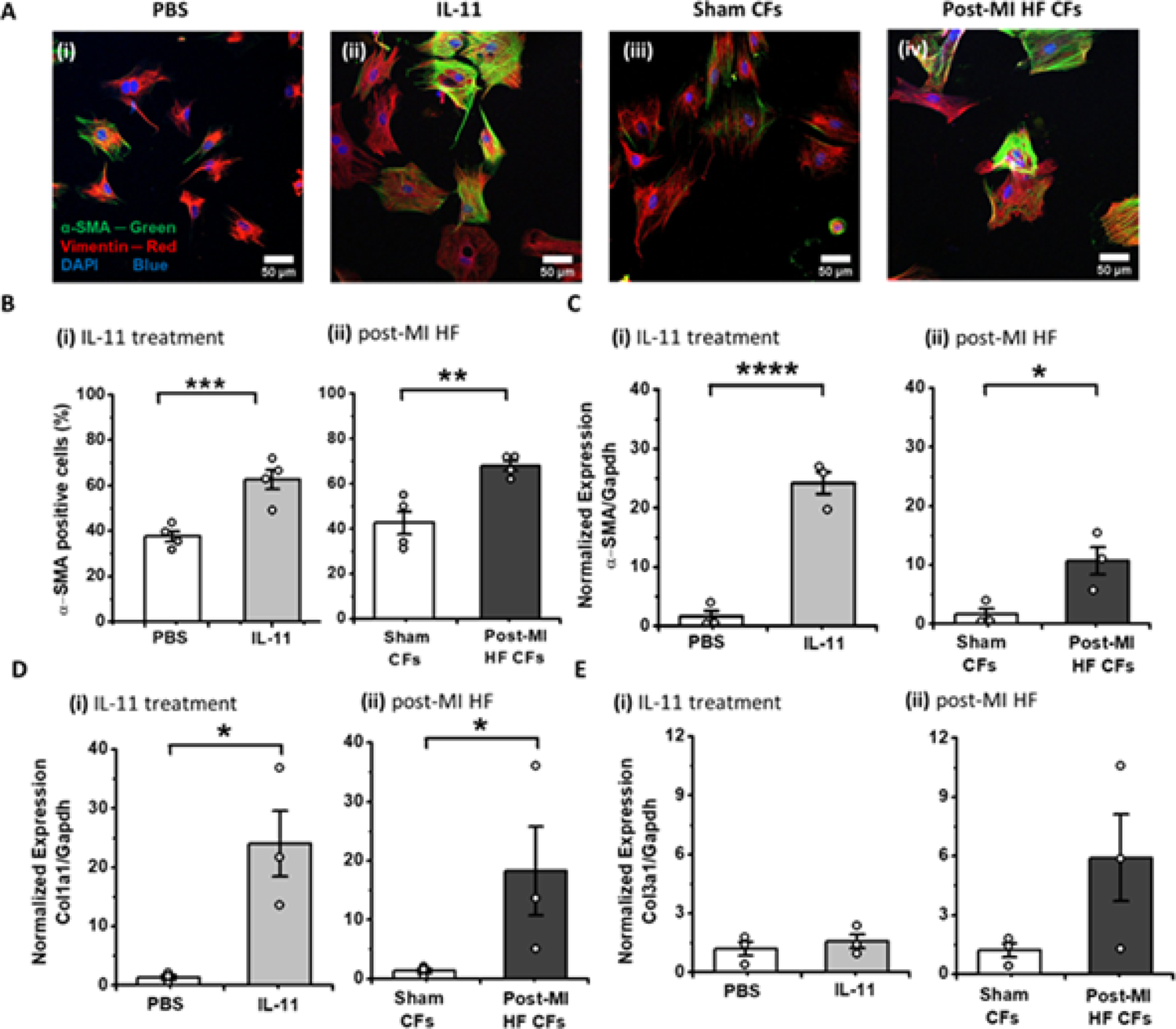
Characterisation of IL-11 and post-MI HF models of fibrosis by immunostaining and RT-qPCR. **A)** Representative images of rCFs stained for α-SMA (green), vimentin (red) and DAPI (blue): rCFs i) treated with PBS; ii) treated with IL-11 (5 ng/µl, 24hrs), iii) harvested from the matched sham-operated rats, iv) derived from post-MI HF rat; **B)**-**E)** experiments realized for the two protocols: **i)** in vitro stimulation with IL-11 (vs PBS) and **ii)** harvesting from the failing and sham-operated hearts. **B)** Quantification of α-SMA positive cells (%) (N = 4, n = 2). ∼200 cells were quantified for each condition, with 10 cell/image. RT-qPCR analysis showing **C)** α-SMA, **D)** Col1a1 and **E)** Col3a1 mRNA expression (normalized to GAPDH) (N = 3; n =3). Data presented as mean fold change values ± SEM in a linear scale. Mann-Whitney test was used to determine significant differences between groups. *p<0.05, **p<0.01, ****p<0.001.

### miRNA-497-5p and miRNA-27b-5p are downstream effectors of IL-11

Bioinformatic target prediction analyses (Supplementary Figure 3) lead to the identification of 68 miRNAs candidates as modulators of the profibrotic responses induced by IL-11 and TGFβ1. Using the criteria described in method and summarized in (Figure 2A), we next filtered for 7 robust miRNA candidates (Figure 2B), which were next investigated by miRNA PCR analysis in IL-11-treated rCFs and in rCFs derived from the failing hearts vs age-matched controls. Preliminary estimation of Ct values demonstrated that all 7 miRNAs were detectable in rCFs. Furthermore, miRNA-27b-5p (Figure 2B-i), miRNA-497-5p (Figure 2B-ii), and miRNA-21-5p (Figure 2B-iii), emerged to be the only miRNAs up-regulated under both profibrotic conditions. Since miRNA-21-5p has been already widely studied in the context of cardiac fibrosis [18], we focused our study on the remaining miRNA-27b-5p and miRNA-497-5p.

**Figure 2.**
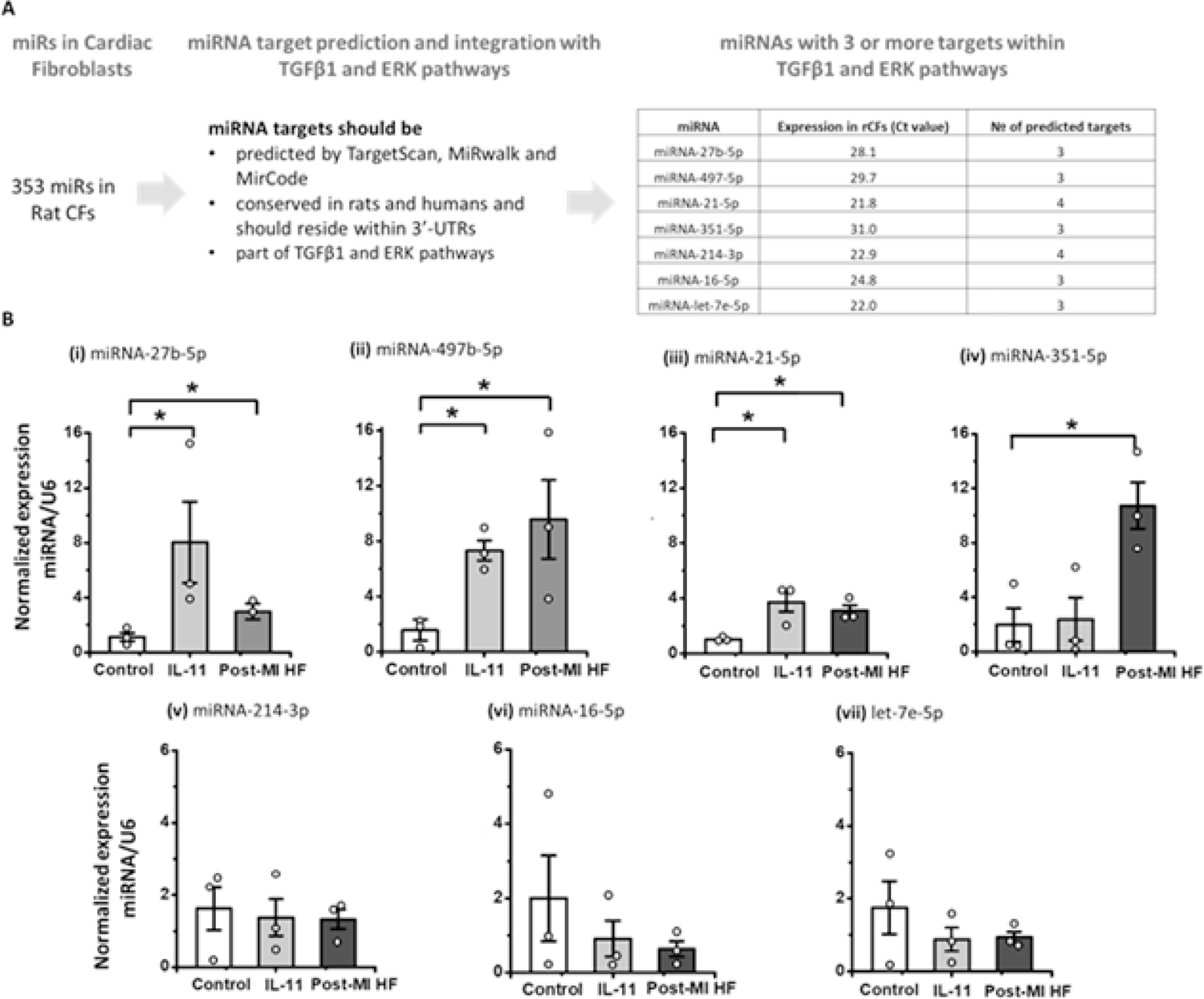
Prediction and validation experiments for miRNAs expressed in cardiac fibroblasts and associated with cardiac fibrosis through regulation of IL-11 and TGFβ1 pathways. **A)** 353 Predicted candidates for identifying miRNAs that are associated with genes obtained from KEGG for the IL-11 (ERK) and TGFβ1 profibrotic pathways. 7 miRNAs were predicted to mediate fibrosis upon IL-11 stimulation. **B)** RT-qPCRs of miRNA expression normalized to U6 in control rCFs (white bar), rCFs treated with IL-11 (light grey) and in post-MI HF rCFs (grey): **i)** miRNA-27b-5p, **ii)** miRNA-497-5p, **iii)** miRNA-21-5p, **iv)** miRNA-351-5p, **v)** miRNA-214-3p, **vi)** miRNA-16-5p, **vii)** miRNA-let-7e-5p. Data are presented as mean fold change values ± SEM in a linear scale (N =3, n =3). One-Way-ANOVA with Tukey’s post-hoc test was used to determine significance between groups. *p<0.05.

### miRNA-27b-5p and miRNA-497-5p are pro-fibrotic in rat cardiac fibroblasts

To evaluate the functional role and significance of miRNA-27b-5p and miRNA-497-5p, we overexpressed or inhibited their expression in rCFs isolated form healthy hearts. Forced expression of either of the two miRNAs increased the percentage of α-SMA positive cells (Figure 3A-i-v). These observations were further confirmed by quantifying the number of a-SMA positive cells (Figure 3B-i-ii). We confirmed that the administration of miRNAs mimics and inhibitors to healthy rCFs successfully altered the gene expression of the corresponding miRNA (Figure 3C,i-ii). Transfection of mimics for each miRNA significantly increased the expression of α-SMA and Collagen 1 (Figure 3D, E). Incubation of cells with inhibitors, significantly reduced Collagen 1 expression but not α-SMA (Figure 3D, E).

**Figure 3.**
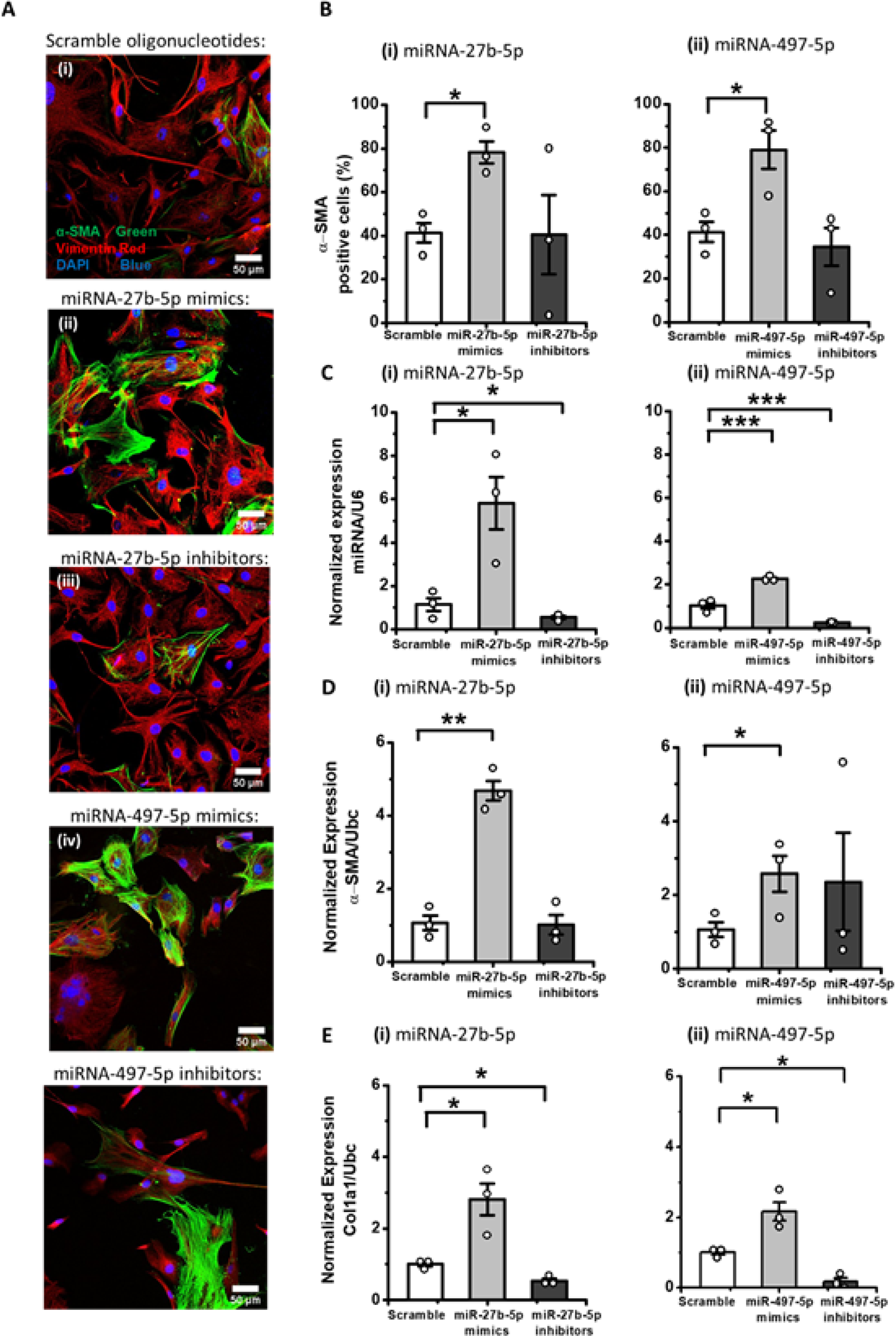
miRNA-27b-5p and miRNA-497-5p induce pro-fibrotic changes in cardiac fibroblasts. Transfection of healthy rCFs with scramble nucleotides (white bar), miRNA-27b-5p or −497 mimics (light grey) and inhibitors (dark grey) at 5nM concentration. **A)** Representative images i)-v) stained after 24h with vimentin (red) and α-SMA (green) and DAPI (blue). **B)-E)** experiments realized for the two gain/loss of functions protocols: **i)** miRNA-27b-5p and **ii)** miRNA-497-5p. **B)** Quantification of α-SMA positive cells (%) (N = 3, n = 2). **C)-E)** PCR analysis of miRNA/mRNA of interest: expression was normalized to U6 RNA expression and further referred to the scramble condition, while mRNA expression was normalized to UBC RNA expression. Expression of **C)** miRNA-27b-5p and miRNA-497-5p, **D)** α-SMA and **E)** Col1a1. Values are represented as mean fold change values ± SEM in a linear scale (N = 3, n = 3). One-way ANOVA was applied followed by Tukey’s post-hoc test was performed in order to determine significance *P-value<0.05, **p<0.01, ***p<0.005.

Next, we checked how miRNA overexpression and inhibition affected rCFs derived from failing hearts. A small but significant boost of α-SMA positive cells driven by miRNA mimics was observed *in vitro* (Figure 4A, B). This effect was accompanied by regulation of the relevant miRNA expression in gain and loss of function experiments (Figure 4 C). Unlike our previous experiment conducted in healthy rCFs, transfection of miRNA inhibitors in post-MI HF rCFs decreased expression of α-SMA, only (Figure 4 D, E).

**Figure 4.**
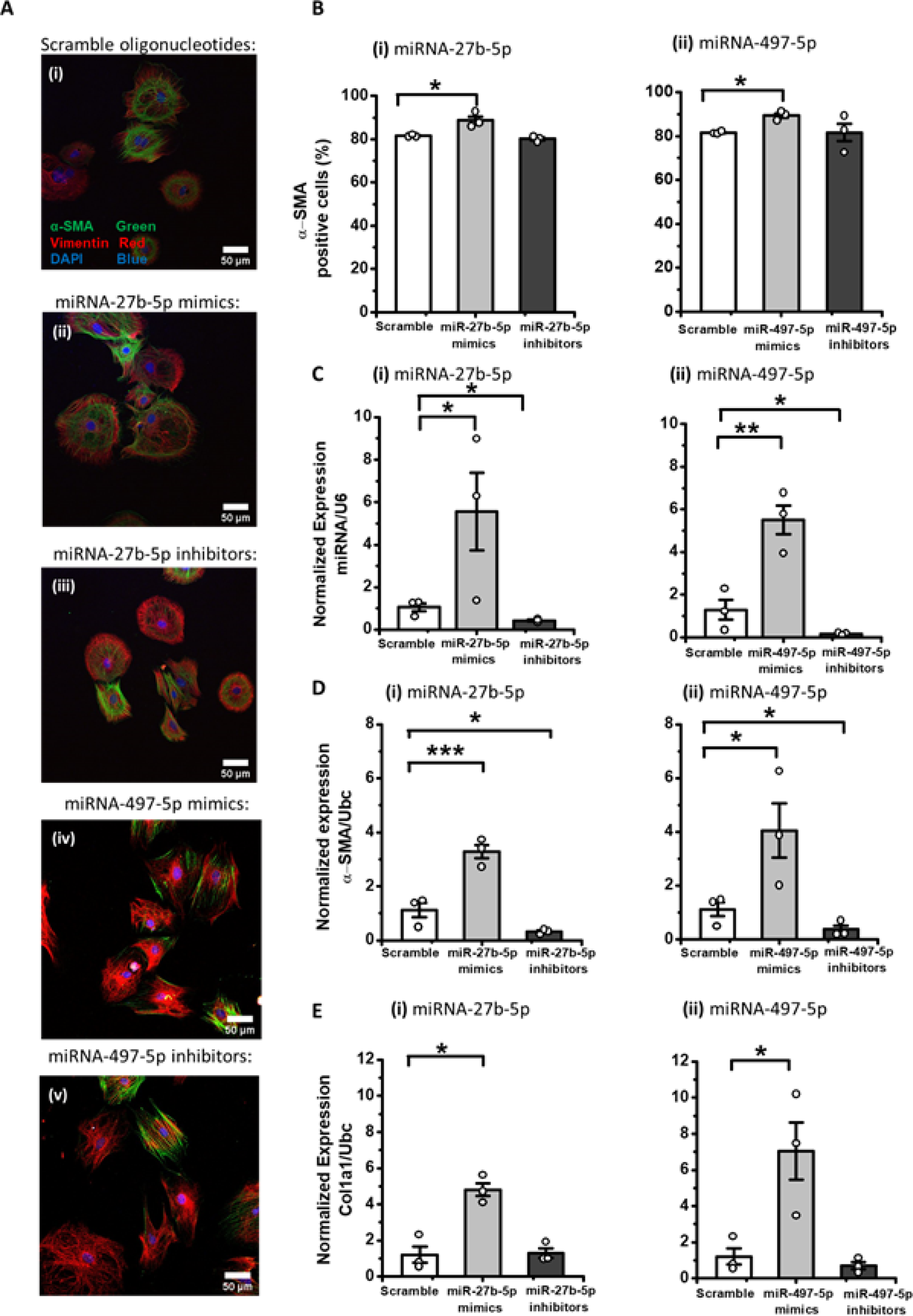
miRNA-27b-5p and miRNA-497-5p modulation induces pro-fibrotic changes in cardiac fibroblasts from failing hearts. Transfection of post-MI HF rCFs with scramble nucleotides (white bar), miRNA-27b-5p or −497 mimics (light grey) and inhibitors (dark grey) at 5nM concentration. **A)** Representative images i)-v) stained after 24h with vimentin (red) and α-SMA (green) and DAPI (blue). **B)-E)** experiments realized for the gain/loss of functions protocols: **i)** miRNA-27b-5p and **ii)** miRNA-497-5p. **B)** Quantification of α-SMA positive cells (%) (N = 3, n = 2). **C)-E)** PCR analysis of miRNA/mRNA of interest: miRNA expression was normalized to U6 RNA expression and further referred to the scramble condition, while mRNA expression was normalized to UBC RNA expression. Expression of **C)** miRNA-27b-5p and miRNA-497-5p, **D)** α-SMA and **E)** Col1a1. Values are represented as mean fold change values ± SEM in a linear scale (N = 3, n = 3). One-way ANOVA was applied followed by Tukey’s post-hoc test was performed in order to determine significance *P-value<0.05, **p<0.01, ***p<0.005.

### miRNA-27b-5p and miRNA-497-5p converge to regulate hypoxia-inducible factor 1 signalling (HIF-1) by targeting EGLN

As described in methods, for miRNA-27b-5p and miRNA-497-5p we predicted 596 and 140 potential gene targets, respectively. 254 predicted targets were found to be differentially expressed in hCFs stimulated with IL-11 as compared to controls. Consequently, they were used for a pathways enrichment test (Figure 5A-i). As a result, the HIF-1 signalling pathway was identified to be enriched with potential targets of miRNA-27b-5p and miRNA-497-5p (Figure 5A-ii). Interestingly, both miRNAs (in yellow) were predicted to target the EGLN genes, which are (Figure 5A-iii, in blue).

**Figure 5.**
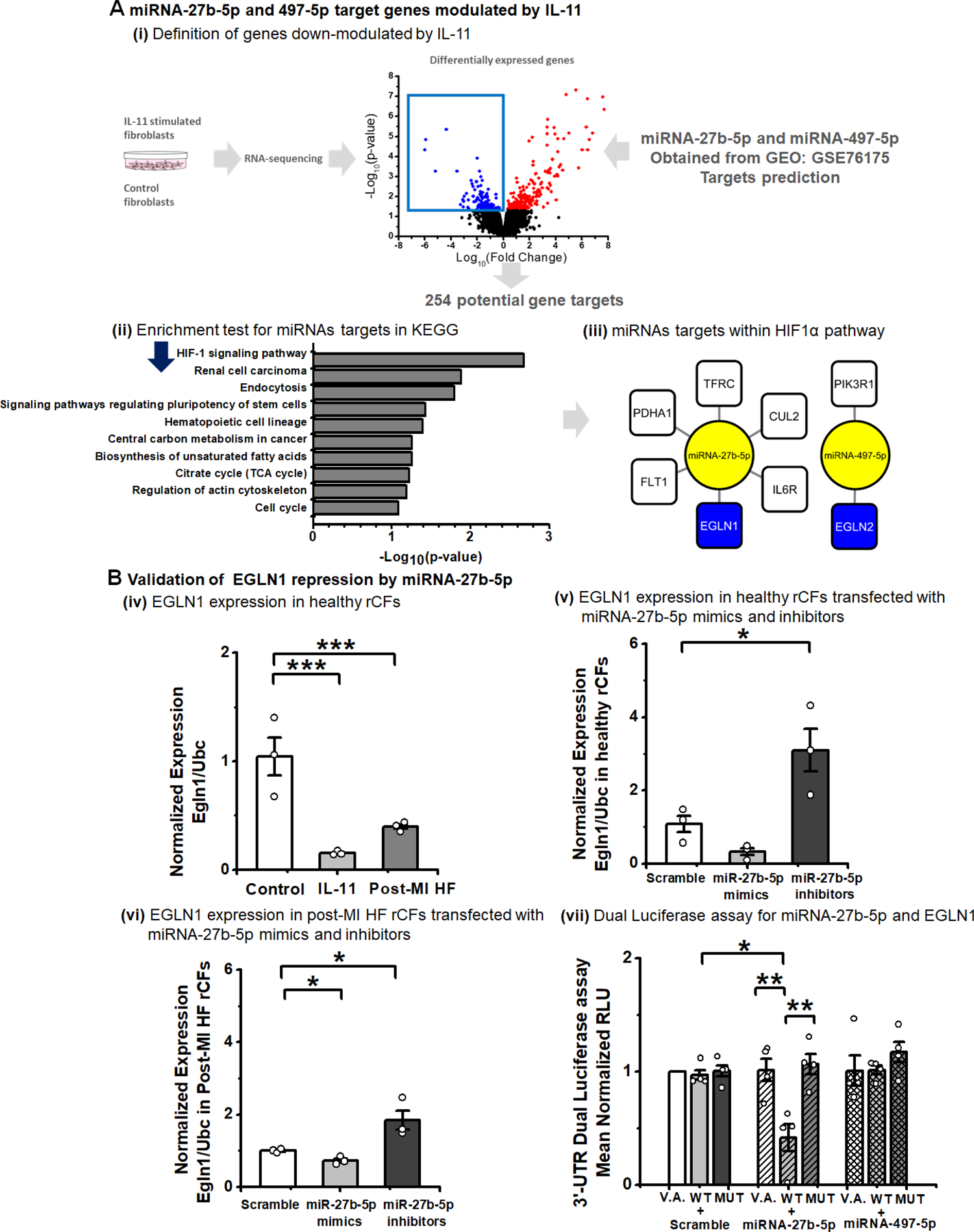
Identification, prioritization and validation of EGLN1 as miRNA-27b-5p target. **A)** Schematic for the description of the target prediction process by using **(i)** down-regulated genes in hCFs stimulated with IL-11 and dataset GEO: GSE76175. **(ii)** Enrichment test: Graph is plotted as – Log10(p-values), p-values<0.05. **(iii)** miRNA targets network from HIF-1 signaling pathway. miRNAs ─ yellow, EGLN1,2 ─ blue, remaining targets ─ white. **B)** Validation of EGLN1 repression by miRNA-27b-5p was performed by PCR analysis of EGLN1 expression in **i)** CFs from healthy rats treated with IL-11 (5ng/µl, 24hrs) (light grey) and post-MI HF CFs (grey), **ii)** healthy rat CFs **iii)** and post-MI HF CFs transfected with 5nM scramble (white bar), miR-27b-5p mimic (light grey) or inhibitor (dark grey). **iv)** 3’-UTR Dual luciferase assay in HEK293FT cells done with a combination of vector alone (V.A.), miR-27b-5p binding site (WT), or a mutated miR-27b-5p binding site (MUT) for EGLN. pLUC plasmid was co-transfected with a plasmid encoding Renilla luciferase along with a plasmid encoding either miR-27b-5p; scramble sequence (scr); miR-497-5p as a negative control. Firefly luciferase luminescence was normalized against renilla luminescence. Experimental values were compared to the vector-only and scramble vector transfection conditions. N = 3 for RT-qPCR. N = 4 for dual luciferase assay. Data presented as Mean ± SEM. One-way ANOVA was applied followed by Tukey post-hoc test to determine statistical significance *P-value<0.05, **P-value<0.01, ***P-value<0.005.

The EGLN (egl nine homolog) gene encodes prolyl hydroxylase domain (PHD) enzymes, which target hypoxia-inducible factor (HIF)-α signalling its polyubiquitination and proteasomal degradation [42]. EGLN plays a critical role in regulating HIF abundance and oxygen homeostasis [42]. The EGLN1 gene/PHD2 protein was recently shown to be an important player in cardiac fibrosis and hypertrophy [43]. Thus, we hypothesised that miRNA-27b-5p targets EGLN1. On the other hand, miRNA-497-5p targets EGLN2, which has already been validated as a direct target of miRNA-497-5p [44]. We therefore embarked to validate the direct binding and expressional regulation of EGLN1 by miRNA-27b-5p. EGLN2 was measured in clinical samples, only (see below).

EGLN1 was significantly down-regulated in IL-11 treated rCFs and post-MI HF rCFs compared to healthy rCFs (Figure 5B-i). Next, EGLN1 expression was tested in healthy and post-MI HF rCFs transfected with miRNA-27b-5p mimics and inhibitors. In both healthy (Figure 5B-ii) and pathological CFs (Figure 5B-iii), miRNA-27b-5p loss of function lead to an increased EGLN1 expression. In post-MI HF CFs, miRNA-27b-5p overexpression resulted in EGLN1 down-regulation. To further validate, EGLN1 direct targeting by miRNA-27b-5p, we conducted a dual luciferase assay (Figure 5B-iv). Our results demonstrated a significant decrease in relative luminescence when miRNA-27b was co-transfected with p-MIR luciferase reporter with the WT EGLN1 binding site (WT EGLN1), but not in the presence of the mutated binding site (MUT EGLN1). Therefore, miRNA-27b-5p showed an affinity for EGLN1. In addition, the pro-fibrotic role of miRNA-27b-5p in rat CFs (Figure 3-5) was confirmed in human CFs (Supplementary 4). Interestingly, along with the decrease in expression of EGLN1 (Supplementary Figure 4C), we observed a significant up-regulation of HIF2α, a down-stream effector of EGLN1 (Supplementary 4D).

### Characterisation of the potential of miRNA-27b-5p and miRNA-497-5p as circulating biomarkers of cardiac fibrosis

LV samples were collected from patients undergoing concomitant septal myectomy during surgical AVR and underwent histological analysis. In these samples there was extensive replacement and interstitial fibrosis when compared to the control group (Figure 6A-i). Quantification of the collagen area (Figure 6A-ii) demonstrated an increase of ECM area in AS patients. This observation was supported by immunostaining against Collagen I (the main collagen type in the ECM) (Figure 6B-i). Quantification on Collagen I area in these biopsies revealed a significant fibrosis increase in AS patients (Figure 6B-ii). In addition, we measured CTGF expression in snap frozen LV biopsies and control group patients. CTGF is a matricellular protein which is actively produced in injured cardiomyocytes and cardiac fibroblasts [45]. We observed significant up-regulation of CTGF expression in LV of AS patients (Figure 6C-i).

**Figure 6.**
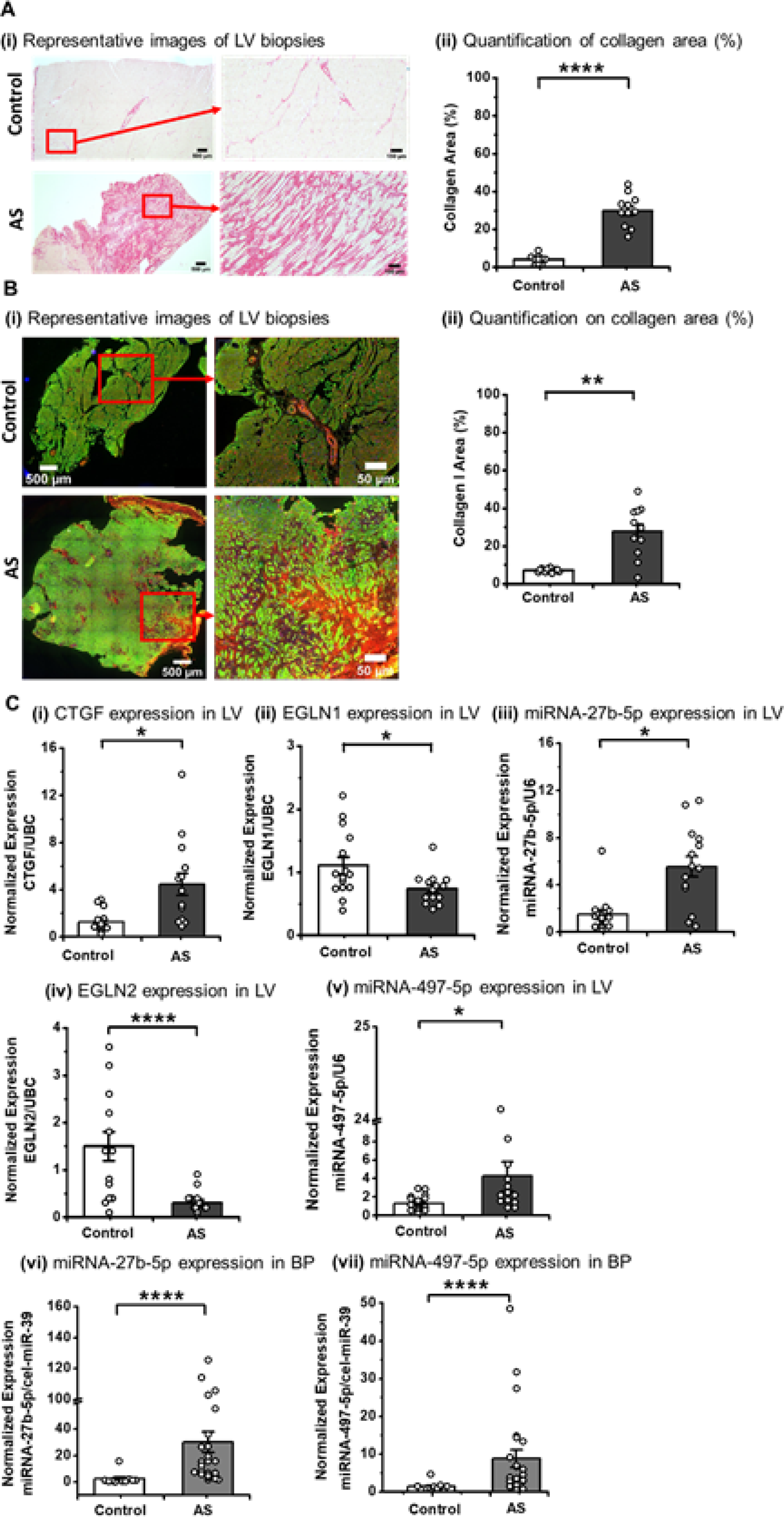
miRNA-27b-5p and miRNA-497-5p are up-regulated in LV and plasma of AS patients characterised with increased collagen deposition. **A)** Histological analysis of LV biopsies from a control and AS groups contains **i)** representative images of LV biopsies from control group and AS patients. Biopsies were stained with sirius red. **ii)** Quantification on collagen area (%) in LV biopsies stained with sirius red in control (white bar) N = 10 and AS group (dark grey) N = 11. **B)** Collagen I staining LV biopsies from a control and AS groups contains **i)** representative images on LV biopsies from patients in control group and AS patients. Biopsies were stained with against collagen I (red), α-SRC (green) and DAPI (blue); **ii)** quantification on collagen I area in stained LV biopsies. **C)** PCR analysis of gene expression normalized to UBC for mRNA and to u6 for miRNAs in LV. miRNAs expression in plasma was normalized to pre-added spike-in (cel-miR-39-5p). Gene expression in LV of AS (dark grey) N = 14 and control (white) patient N = 14 of **i)** CTGF, **ii)** EGLN1, **iii)** miRNA-27b-5p **iv)** EGLN2, **v)** miRNA-497-5p. miRNAs expression: **vi)** miRNA-27b-5p, **vii)** miRNA-497-5p in plasma of AS (grey) N = 24 referred to a control group (N = 10). Values presented as mean ± SEM in a linear scale. Mann Whitney test was performed in order to estimate statistical significance*P-value<0.05. **p-value<0.01. ***p-value<0.005. ****p-value<0.001.

Next, we measured the expression of the miRNA-27b-5p target EGLN (Figure 6C-ii;). We observed a significant decrease of EGLN1 expression in the LV of AS patients. These data are in keeping with the up-regulation of miRNA-27b-5p in the same biopsies (Figure 6C-iii), which was also accompanied by increased expression of miRNA-497-5p and supressed expression of EGLN2 (Figure 6C-iv,v).

As a further step, we measured the expression of both miRNA-27b-5p and miRNA-497-5p in the peripheral blood plasma of AS patients, comparing their expression with sex matched healthy donors. Interestingly, the levels of both miRNAs were increased in plasma of AS patients (Figure 6C-vi,vii). As described in table 1, there were significant differences in the AS and control groups in terms of age, presence of hypertension and dyslipidaemia. For this reason, we performed a sub-analysis on miRNA-27b-5p, miRNA-497-5p and EGLN1 expression in LV of AS patients with hypertension or dyslipidaemia (Supplementary 5). We demonstrated that measured values are not different in terms of expression between AS patients with or without hypertension (Supplementary Figure 5A-C). No significant regulation of miRNAs and EGLN1 expression was observed in dyslipidaemia settings (Supplementary figure 5D-F). In addition, miRNA-27b-5p, miRNA-497-5p and EGLN1 expression in LV of AS patients did not significantly correlate with patient age.

miRNAs can circulate in the blood encapsulated in extracellular vesicles (EVs) [22,46]. We isolated EVs from plasma of AS patients and healthy donors. EVs (50-130 nm diameter) were characterised by transmission electron microscopy with negative uranyl acetate staining (Figure 7A). As it was observed in the plasma, miRNA-27b-5 (Figure 7B-i) and miRNA-497-5p (Figure 7B-ii) level was significantly higher in EVs derived from AS patients.

**Figure 7.**
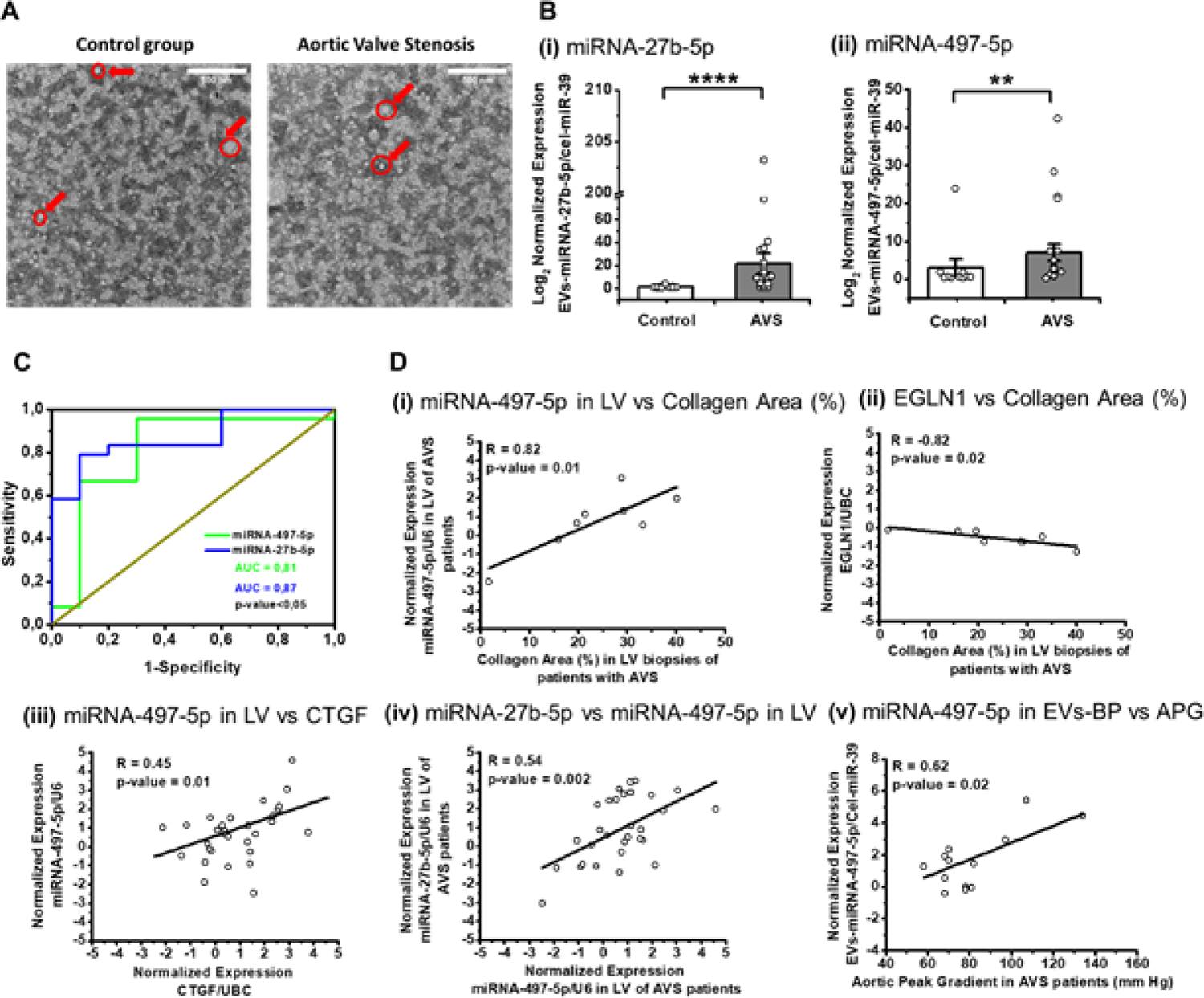
miRNA-27b-5p and miRNA-497-5p expression in EVs from plasma and correlations with expression of fibrotic markers. **A)** Representative images of EVs isolated from plasma. Red arrows and circles highlight single EVs. **B)** Expression of **i)** miRNA-27b-5p and **ii)** miRNA-497-5p in EVs from plasma (N = 23) (N = 10). Statistical significance was evaluated with Mann-Whitney test **p-value<0.01, ****p-value<0.001. **C)** ROC curve analysis based on expression of miRNA-497-5p (green), miRNA-27b-5p (blue) in EVs from plasma of AS (N = 24) and healthy donors (N = 10). Area under the curve is measure and present on the graph along with p-values. **D)** Pearson correlations on measured parameters in AS patients between: **i)** normalized expression of miRNA-497-5p in LV of AS patients with Collagen Area (%) measured in LV biopsies stained with sirius red (N = 8); **ii)** mean normalized expression of EGLN1 in LV with Collagen Area (%) measured in LV stained with sirius red (N = 8); **iii)** normalized expression of miRNA-497-5p in LV with normalized expression with CTGF in LV (N = 29); **iv)** normalized expression of miRNA-27b-5p in LV with normalized expression of miRNA-497-5p in LV (N = 29); **v)** normalized expression of miRNA-497-5p in EVs from plasma of AS patients with patients aortic peak gradient (mmHg) (N = 14).

ROC Curves were plotted based on miRNA levels in EVs isolated from plasma (Figure 7C). Both miRNA-27b-5p and miRNA-497-5p discriminated AS patients from controls in a sensitive and specific manner, as indicated by the significantly high area under the curve (AUC) for each miRNA. Following this, we tested correlations between miRNA expression in LV, plasma and EVs from plasma with histology data, CTGF expression and clinical parameters in patients with a Pearson’s correlation test. We found that miRNA-497-5p expression in LV positively correlated (R=0.82, p-value = 0.01) with collagen area (%) (Figure 7D-i), CTGF expression (R = 0.45, p-value = 0.01) (Figure 7D-iii) and with miRNA-27b-5p expression in LV (R = 0.54, p-value = 0.002) (Figure 7D-iv). At the same time, the miRNA-27b-5p target, EGLN1, was negatively correlated (R = −0.82, p-value = 0.02) with collagen area (%) (Figure 7D-ii). Surprisingly, miRNA expression in plasma did not show any correlation with clinical parameters. Interestingly, miRNA-497-5p levels in EVs derived from plasma of AS patients was positively correlated (R = 0.62, p-value = 0.02) with aortic peak gradient in AS patients.

In addition, we tested how risk factors such as age, hypertension and dyslipidaemia affected miRNA-27b-5p and miRNA-497-5p-5 expression in plasma and EVs derived from plasma (Supplementary 6). We found that these factors did not affect miRNAs expression. However, miRNA-497-5p expression in EVs derived from plasma demonstrated some dependence on patient age (Supplementary 6L).

## Discussion

The results of our study candidate miRNA-27b-5p and miRNA-497-5p as IL-11-responsive mediators of cardiac fibrosis. Moreover, we have additionally identified the potential of the two miRNAs as circulating biomarkers of cardiac fibrosis in AS patients with preserved ejection fraction.

We report that the profibrotic IL-11 increases miRNA-27b-5p and miRNA-497-5p in healthy CF and the 2 miRNAs are also increased in post-MI HF CFs. Previous studies suggested a cardio-protective and anti-fibrotic effects of IL-11 through JAK-STAT signalling in the mouse heart in the context of ischemia-reperfusion injury [7]. However, recently this concept has been reconsidered by Schafer *et al.* [6], indicating a deep need of further investigation of IL-11 non-canonical molecular mechanisms, such as through regulations of miRNAs. miRNA-27b-5p and miRNA-497-5p were previously investigated in the fibrosis settings [47–49]. In detail, miRNA-27b enhanced cardiac fibrosis [47], and miRNA-497-5p regulates ECM remodelling in the lungs [48,49].

Our data also highlights a potential mechanism of fibrosis induction by miRNAs through expressional repression of EGLN, a family of genes encoding PHD enzymes that regulate the stability of HIFs. HIFs are hypoxia inducible transcription factors which are also important in pro-fibrotic gene expression [50,51]. HIF comprises two subunits: the constitutively expressed HIF-β and HIF-α, with a short half-life under homeostatic conditions. PHD-induced hydroxylation triggers the degradation of HIF-α by allowing the binding of the von Hippel-Lindau tumour (pVHL) suppressor, which targets HIFα for proteasomal degradation [52]. Both hypoxia and EGLN repression improve HIFα stabilization and facilitates a myriad of HIF-controlled transcriptional responses. While short-term HIF-1α stabilization has been reported to have a cardioprotective impact, the long-term HIF-1α activation drives autonomous pathways that add to disease progression; accordingly, cardiomyocyte-selective HIF-1α transgenic mice develop heart failure. Moreover, EGLN/PHD loss has been functionally implicated with cardiac fibrosis and hypertrophy [43,53]. Recently, Dai *et al* demonstrated that mice with endothelial-specific *EGLN1* gene deletion exhibited left ventricle hypertrophy and cardiac fibrosis and identified HIF2α as a transcriptional factor responsible for the pro-fibrotic signalling [43]. Interestingly, IL-11 differential co-expression network is enriched in genes associated with hypoxia (e.g., HIF2α, VEGFC, CITED2) [37]. These findings are in keeping with previous reports of EGLN2 to be a target of miRNA-497-5p [44], and with our demonstration that miRNA-27b-5p directly binds to and suppresses EGLN1 mRNA expression. Reduced EGLN1 expression was also observed in rat CFs treated with IL-11, in CFs extracted from the post-MI failing hearts, and in the LV of AS patients together with EGLN2, supporting the importance of EGLN regulation during the development of cardiac fibrosis in a clinically relevant setting. Moreover, we found that HIF2α expression in human CF treatment increases following miRNA-27b-5p mimic. Thus, IL-11-induced miRNA-27b-5p and miR-497-5p may drive an excessive and profibrogenic hypoxia-signalling *via* regulating both PHD1/ELGN1 and PHD2/ELGN2 (Figure 8). Noteworthy, the disease models studied here are associated with hypoxia. Ischemia in cardiac tissue leads to massive necrosis, which promotes inflammatory cell intervention, fibroblast transdifferentiation and tissue hypoxia [54]. Moreover, the NF-κB-HIF-2 pathway plays a role in AS, where HIF2α promotes Collagen X expression [55].

**Figure 8.**
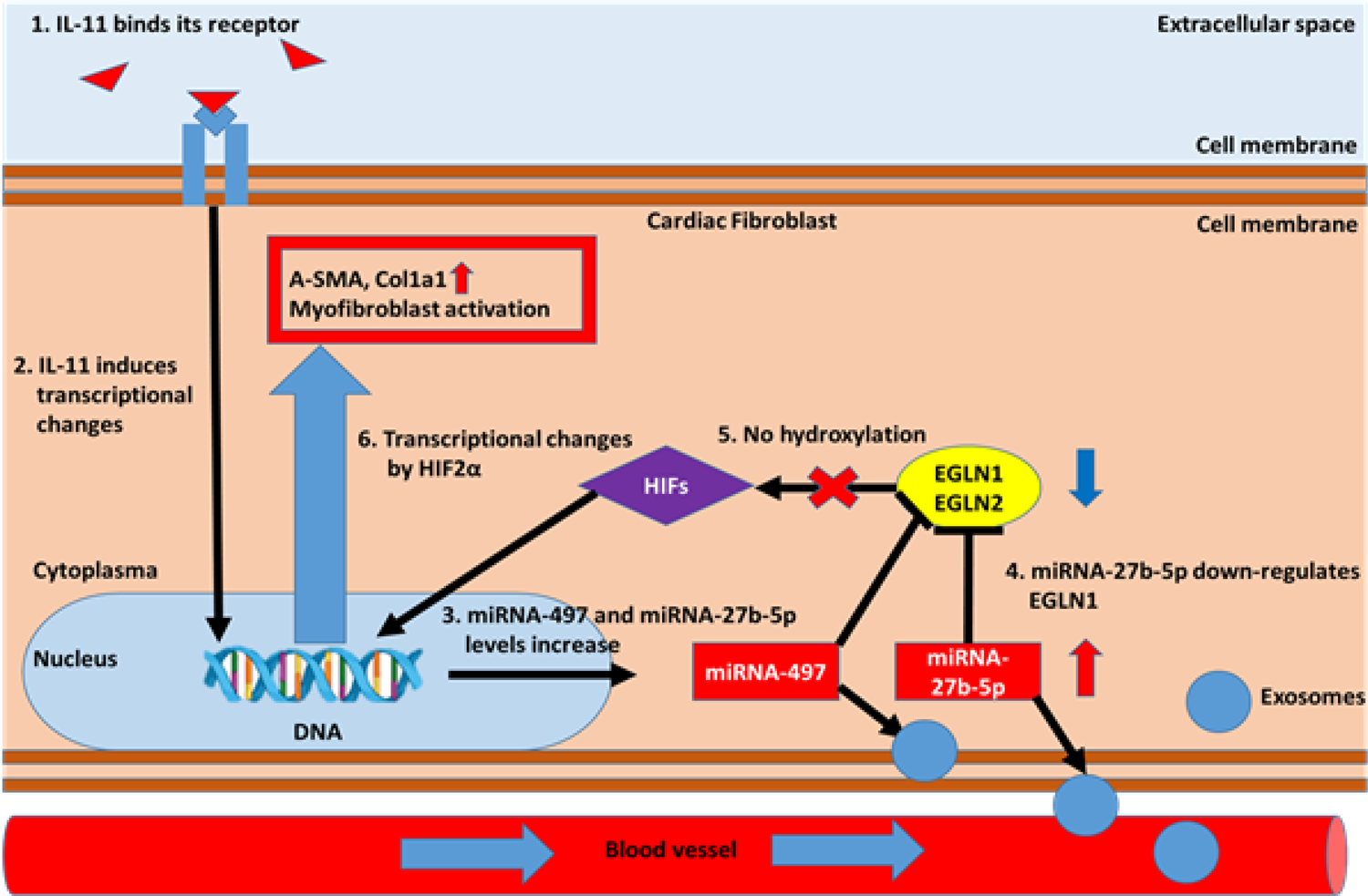
Proposed molecular mechanism of miRNA-27b-5p and miRNA-497-5p contribution in cardiac fibrosis. IL-11 stimulation of CFs leads to increase of miRNA-497-5p and miRNA-27b-5p expression. Following this, miRNA-27b-5p is responsible for EGLN1 suppression (miRNA-497-5p for EGLN2 suppression) and further HIF stabilization and pro-fibrotic transcriptional changes.

Fibrosis represents a key feature of myocardial damage induced by AS and represents the transition between the compensatory hypertrophy with interstitial fibrosis at the early stages and heart failure associated with replacement fibrosis, that is irreversible [56].

The use of circulating biomarkers has been encouraged in recent guidelines for AS monitoring and management [57]. Studying the relationship between BNP clinical activation and long-term outcomes in AS patients previously demonstrated a clear correlation between BNP active levels and mortality, even in asymptomatic patients [58]. However, a threefold increase was associated with the worst survival rate, possibly due to a clinically established HF syndrome. Specific biomarkers associated with fibrosis in AS would identify patients more precisely.

Beyond their potential as therapeutic targets, miRNA-27-5p and miR-497-5p also displayed a potential as clinical-relevant biomarkers of early myocardial fibrosis. We observed an increase in expression of circulating miRNA-27-5p and miR-497-5p in AS patients with preserved ejection fraction. We demonstrated that both miRNA-27b-5p and miRNA-497-5p are up-regulated in the diseased myocardium, plasma and EVs derived from plasma of AS patients characterised with intensive ECM accumulation and LVEF > 55%. Interestingly, both EGLNs, targets of miRNA-27b-5p and miRNA-497-5p, were down-regulated in the myocardium of these patients. Correlation analysis indicated strong correlation of miRNA-497-5p levels in LV with CTGF expression and collagen area in histology slides, while miRNA-27b-5p was positively correlated with size of IVS. At the same time, we showed that miRNAs levels in blood plasma can be potentially used to distinguish healthy patients from AS patients with preserved ejection fraction. Using these data, we report miRNA-27b-5p and miRNA-497-5p as potential biomarkers of cardiac fibrosis in AS patients with preserved ejection fraction.

Our results suggest that the miRNAs could be useful in (1) patients with symptomatic severe AS the for their risk stratification according to the presence of fibrosis and (2) asymptomatic patients with severe AS with preserved LV function under surveillance to determine time of intervention before an irreversible LV damage occurs. Current ESC/EACTS guidelines recommend the treatment of AS when patients begin to experience symptoms or in case of LV impairment [57]. However, symptoms are difficult to assess in elderly comorbid patients and survival rates decrease dramatically after the onset of symptoms [59]. Moreover, these data are limited to small non-randomised series. In asymptomatic patients, treatment is recommended for patients with preserved LV function and critical or rapid progressing AS and should be considered when LV function is below <55% without another cause. However, the latter condition is associated with an ongoing LV damage already established. Historically the cut-off for LV dysfunction has been considered LVEF <50%. In a large retrospective multicentric study including 1,678 patients with severe AS and preserved LV function, LV function below 55% was associated with poor outcomes, despite treatment [60], emphasizing the importance of early recognition of myocardial damage.

Extra-aortic cardiac damage assessment has been proven to be relevant for risk stratification in patients with AS, with evidence of important prognostic implications. Pooled data from the intermediate and high-risk patients of the PARTNER1, 2 and S3 trials have shown increased mortality and hospitalisations at 5 years in case of no LV remodelling regression [61]. Echocardiographic study on the pooled population of the PARTNER2 and 3 trials, have demonstrated that the two-year mortality was associated with extent of cardiac damage at baseline. These results were further confirmed in asymptomatic patients with an updated echocardiographic scoring system in which mild LV dysfunction (LVEF<60%), was associated with increased mortality [62]. These data support the concept that myocardial function assessment by only echocardiogram may not be able to detect an early myocardial damage generated by the continuous pressure overload in AS.

### Limitations

Several studies have demonstrated the importance of identification of myocardial fibrosis on CMR. In particular, the presence of focal scar detected with LGE and its extent were significantly correlated with mortality [56, 62–63]. The EVoLVeD trial (Early Valve Replacement Guided by Biomarkers of LV Decompensation in Asymptomatic Patients with Severe AS), which is actively recruiting, is studying the management of asymptomatic patients with early AVR can improve the adverse prognosis associated with midwall LGE [56]. However, CMR is an expensive, time-consuming resource, not available in all centres and not suitable for all AS patients due to the limitations associated with this technique namely claustrophobia and respiratory artefacts [60]. Thus, CMR is not routinely performed in patients with AS. For these reasons, the identification of circulating biomarkers of fibrosis associated with aortic stenosis, could provide a cost-effective, and time-effective solution to assess fibrosis. A future longitudinal study, associating CMR study of fibrosis and specific sequences to characterise the LV remodelling with levels of circulating miRNA-27-5p and miRNA-497-5p, will give us further insight regarding the dynamics of miRNA, with fibrosis onset, potential fibrosis regression and outcomes in this population. The extremely selected population included in our study, which excludes the other possible cardiac and non-cardiac causes of LV fibrosis highly correlates our findings to LV fibrosis secondary to AS. This could limit the study of miRNA-27-5p and miRNA-497-5p only to a highly selected population. Future studies are needed to validate the study of these miRNAs in a larger cohort of patients with concomitant comorbidities to assess specificity.

### Clinical perspectives

Cardiac fibrosis impairs heart functions and integrity by extracellular matrix (ECM) deposition. In aortic stenosis (AS), fibrosis is one of the key features leading to a decrease in ejection fraction, which is used to make a decision upon surgery for the patients. There is a clinical need to establish reliable blood biomarkers of cardiac fibrosis which can help inform clinical decisions before symptoms present. This is the first study to report on potential utilization of miRNA-497-5p and miRNA-27b-5p as biomarker of cardiac fibrosis in AS patients with preserved ejection fraction. Our study shows the potential of these miRNAs to support the decision-making process for cardiac surgery in AS patients.

### Translational outlook

We found that both miRNAs are critical component of the IL-11 pro-fibrotic mechanism. In particular, we found a novel mechanism which underlies IL-11 signalling in CFs: EGLN1 suppression by miRNA-27b-5p. Thus, this study may serve as a starting point for further investigation upon EGLN1 and hypoxia signalling in cardiac fibrosis.

## Acknowledgements

We are grateful to Maria Carla Panzeri and Valeria Berno and the other staff of the Advanced Light and Electron Microscopy BioImaging Center (ALEMBIC) in San Raffaele Hospital for the technical assistance. We are thankful to Marialucia Longo (IRCCS Policlinico San Donato) for providing human heart tissue sections and to Prof. Cesare Terracciano (Imperial College) for proving human cardiac fibroblasts.

## Abbreviations

LV: Left ventricle

CFs: Cardiac fibroblasts

AS: Aortic stenosis

miRNA: microRNA

IL-11: Interleukin 11

α-SMA: α smooth muscle actin

ECM: extracellular matrix

sEVs: Small extracellular vesicles

post-MI HF: Post myocardial infarction hear failure

EGLN: Egl-9 family hypoxia inducible factor

